# A Practical and Safe Model of Nitrogen Mustard Injury in Cornea

**DOI:** 10.1101/2024.10.18.619116

**Authors:** Ana M. Sandoval-Castellanos, Yao Ke, Tiffany M. Dam, Emanual Maverakis, Mark J. Mannis, Xiao-Jing Wang, Min Zhao

**Author notes:** Corresponding author: Dr. Min Zhao, One Shields Ave. Tupper Hall, Davis, California, 95616 USA. Commercial relationships disclosures: none.

## Abstract

**Purpose:** Sulfur mustard (SM) is an alkylating agent used in warfare and terrorism that inflicts devastating ocular injuries. Although the clinical symptoms are well described, the underlying mechanisms are not fully understood, hindering the development of effective treatments. One major roadblock is the lack of a suitable model due to the extremely hazardous nature of SM, which requires strict safety measures. As a safe and practical alternative, we report a novel model that uses mechlorethamine (nitrogen mustard) gel, an FDA-approved topical chemotherapeutic administered by patients at home. Here we demonstrate its suitability to induce mustard corneal injury in any laboratory.

**Methods:** *Ex vivo* porcine corneas were injured with mechlorethamine gel. Hematoxylin-eosin staining, and immunohistochemistry were performed to evaluate histopathology of SM-like corneal injuries: epithelium thickness and stromal separation, keratocyte and inflammatory cell counts, and expression of inflammation and fibrosis markers.

**Results:** This model showed the characteristic histopathology and expression of cyclooxygenase-2 (inflammation) and fibronectin-1 (fibrosis), which were consistent with other well-established SM-like corneal injury models.

**Conclusion:** Given its ease of implementation and safety, this mechlorethamine model could be used to study the full course of mustard corneal injuries. This model would greatly facilitate mustard injury research, shedding light on new knowledge that would increase our understanding of mustard ocular injuries while investigating novel therapeutics.

**Translational relevance:** this model will allow safe evaluation of SM-like corneal injuries within 24 hours, facilitating the identification of early/new molecules that might help to develop novel treatments which could be readily translated into the clinic.

## 1. Introduction

Sulfur mustard (bis[2-chloroethyl]sulfide; SM) is an alkylating agent used in warfare and terrorism which has inflicted thousands of devastating ocular injuries, affecting the quality of people’s lives ^(1,2)^. SM was first used in WWI at Ypres (1917), and most recently in the Iran-Iraq conflict (1980-1988) and the Syrian Civil War (2011-present) ^(2–5)^. SM is a low-cost and accessible chemical agent, easily synthesized and often stockpiled. Disseminated as vapor, or liquid droplets, SM persists as a threat to soldiers and civilians around the world ^(2)^. The severity of the SM ocular injury varies, depending on dosage and exposure time ^(6,7)^.

Clinically, SM ocular injuries, known as mustard gas keratopathy (MGK), manifest in two phases: an acute phase with symptoms such as eye pain, photophobia, decreased vision, conjunctivitis, and lacrimation; and a chronic phase, where these symptoms reappear alongside new ones, such as neovascularization, edema, corneal opacification, ulceration, dry eye, limbal stem cell deficiency, and even blindness) ^(8–14)^. Current medical treatments include daily irrigation, pain management, anti-inflammatories, antibiotics, limbal stem cell transplantation, and amniotic membrane transplantation, but there is no effective cure for MGK, and hence, suitable therapies are yet to be developed ^(2,5,11,13)^.

The most accepted theory of action is that SM alkylates DNA, damaging not only DNA but also RNA, proteins, and lipid membranes: SM undergoes cyclization and forms ethylene sulfonium, which is later converted to carbonium ions which react with DNA, RNA, and proteins. Cells try to repair DNA by activating poly (ADP-ribose) polymerase (PARP). However, excessive PARP activity causes a reduction of nicotinamide adenine dinucleotide (NAD^+^), decreasing glycolysis. This process hinders energy production, causing cell death ^(11,15–17)^. Additionally, DNA damage causes errors in DNA replication leading to the synthesis of aberrant proteins ^(17)^, that result in abnormal corneal wound healing. Even though angiogenesis, fibrosis, oxidative stress, inflammation (through the production of cyclooxygenase-1 (COX-1) and COX-2)), and expression of fibronectin and matrix metalloproteinases (MMPs) are indicators of corneal SM injury, the biological mechanisms responsible for MGK are poorly understood ^(7,8,10,12–14)^.

Aside from the complexity of the mechanism of action of SM, another roadblock arises while studying SM-induced injuries: SM is an extremely hazardous material, which requires highly controlled research environments and strict regulations and permits that are not available to most ocular research laboratories in the USA ^(18,19)^. SM presents a grave danger to scientists, as accidental exposure can be catastrophic ^(5)^. For this reason, biologically relevant models to investigate the mechanistic effects of SM are scarce ^(13)^. Nitrogen mustard (bis(2-chloroethyl) methylamine, NM) is also an alkylating agent, analogous to SM ^(13,20)^. NM, like SM, modifies DNA, proteins, and other molecules, causing ocular injuries similar to exposure to SM ^(13)^. Despite NM being commercially available, it has the disadvantage that it is a very toxic agent that causes corrosion, acute dermal and ocular toxicity, and which possesses severe mutagenic and carcinogenic properties ^(21)^. In addition, NM reagent is sold as a powder; hence, preparation is needed, increasing the risk of eye, skin, and pulmonary exposure. Therefore, it is imperative to find an agent that can mimic the disastrous effects of SM or NM, without hindering the researchers’ health and safety.

Current mustard injury models, both *ex vivo* and *in vivo*, use vapor SM or liquid NM and study the pathology, histopathology, and molecule expression of diverse biological pathway mechanisms. The animals used in these models are mice, rats, bovines, and rabbits. However, even though corneas from rats and mice and their specific reagents are available, their anatomy differs from human corneas ^(9,12,22)^. The use of rabbit eyes has proven to be advantageous due to their anatomical similarities to the human eye. Nevertheless, rabbit corneas are more resistant to SM injury due mainly to differences in the cornea’s permeability ^(12,23)^. Porcine corneas are emerging as a corneal tissue of choice because they are biologically similar to human corneas, cost-effective, readily available, easy to handle, and follow the 3Rs principle in animal research (replacement, reduction, and refinement) ^(24–26)^.

Our scientific question was whether we could develop an NM-induced cornea injury model using a safer alternative to SM and liquid NM. Therefore, we developed a model using 0.016% mechlorethamine gel, also known as NM. Mechlorethamine gel is currently used for the topical treatment of stage IA and IB mycosis fungoid-type cutaneous T-cell lymphoma ^(27)^. Mechlorethamine gel is safe for patients to use in their homes; hence, it is a safe drug to handle in the laboratory without the need for specialized protective equipment or approved facilities.

Herein we report a safe, novel, and practical NM-induced corneal injury model. We delivered an NM injury, using topical mechlorethamine on *ex vivo* porcine corneas. We observed that changes in epithelial thickness, loss of epithelial layer, (de-epithelialization), epithelial-stroma separation, and decreased keratocyte cell count were present in the corneal tissue. To further validate this model, we evaluated the production of COX-2 and fibronectin 1 (FN1) by immunohistochemistry (IHC), as SM and NM injury induces inflammation and fibrosis. Our results show an increased expression of both markets after NM exposure.

The epithelial histopathology and expression of inflammation and fibrotic markers, shown in this mechlorethamine gel model, are consistent with those in well-established SM- and NM-induced corneal injury models, thus providing a practical and safe model that can be used in any laboratory to study vesicant-induced injuries in the cornea, and for the development of novel therapeutics.

## 2. Materials and methods

### 2.1 Corneal tissue

Porcine eyes (from male and female pigs, 180-300 lbs.) were obtained from Sierra Medical Inc. (Whittier, CA, USA) and the Meat Laboratory at UC Davis (Department of Animal Science, Davis, CA, USA). Eyes were processed immediately after arrival. Excess tissue (fat, muscle, connective) was removed using surgical scissors. Then, globes were rinsed twice with sterile phosphate buffered saline (PBS, Amresco, Cat No. E404-200TABS, USA). For excising the corneas, the protocol by Castro et al. ^(28)^ was followed, with some modifications: the globe was held with a tissue (Kimwipes, Kimberly-Clark Professional, USA), and the cornea was excised from the eyeball using a no. 11 blade by making an incision and cutting ∼ 2 mm from the edge of the cornea, to include the limbus. The cornea was then placed upside down in a Petri dish with PBS. Then, with two pairs of forceps, the cornea was held upside down, forming a cup, and filled with warmed 1% (w/v) agar (Sigma-Aldrich, Cat No. A6686, Germany) with1 mg/mL collagen (PureCol Type I collagen solution (bovine), Advance Biomatrix, Cat No. 5005, USA) solution in DMEM/F12 (Dulbecco’s Modified Eagle Medium/Ham’s F-12, Life Technologies, Cat No. 11330032, USA) to maintain the corneal curvature. When the agar-collagen solution hardened, the cornea was placed right side up in a Petri dish and culture medium added until the limbus was covered, creating an air-liquid interface (see Figure 1.A). Corneas were cultured at 37°C with 5% CO_2_ for a recovery period of 24 h. See Supplementary Table S1 for culture medium composition.

**Figure 1.**
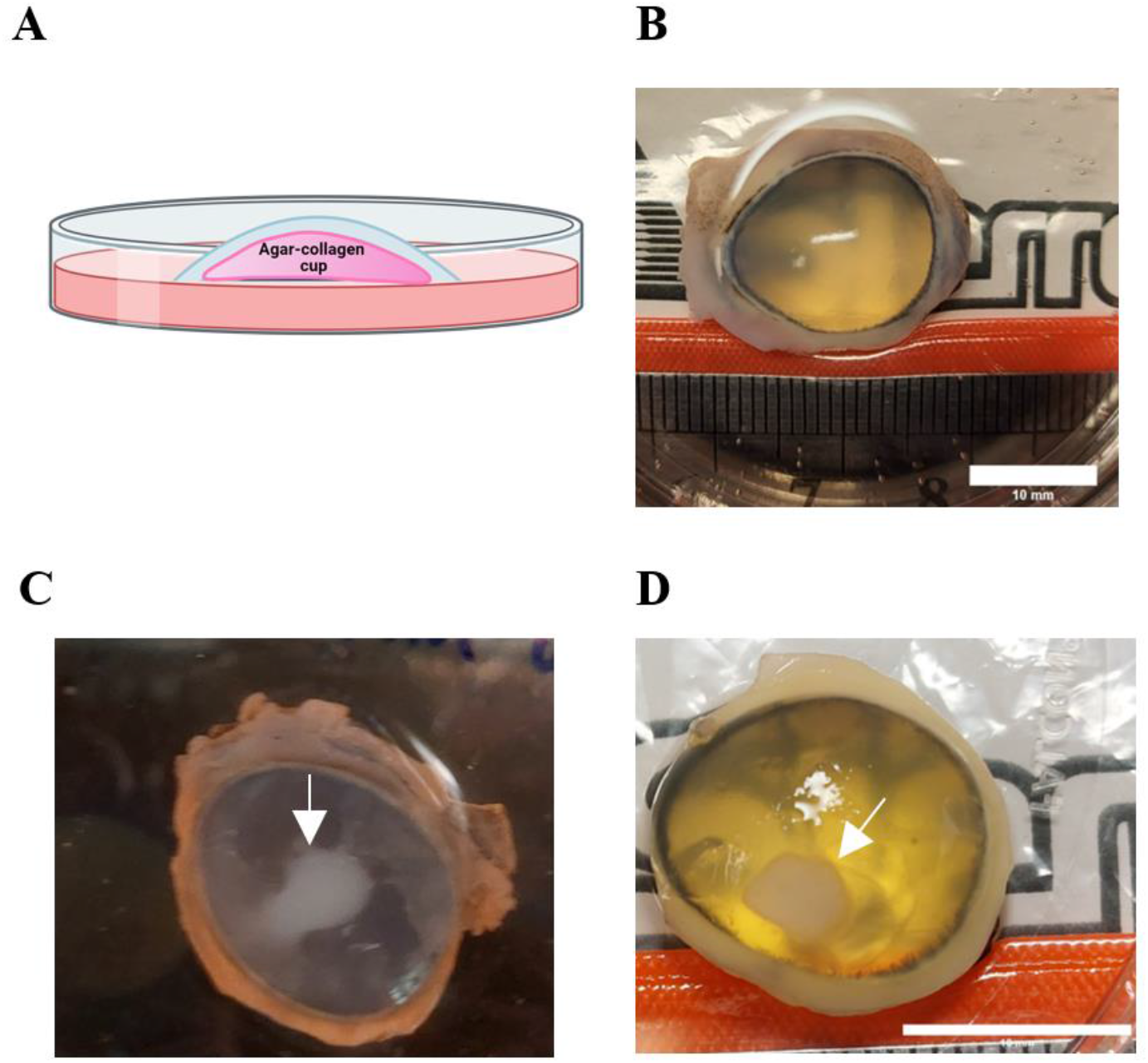
Mechlorethamine gel exposure induces de-epithelialization and opacity of the cornea typically seen in nitrogen mustard (NM)-induced corneal injury. A. Schematic diagram showing the organ culture of the porcine cornea (blue) at the air-liquid interface. B. Organ culture image of a healthy, clear, uninjured porcine cornea, viewed from above. C. and D. A porcine cornea with a NM injury. An opaque area is seen at the site of injury (white arrow). The image was taken immediately after wounding. Scale bars = 10 mm.

### 2.2 Injuring the cornea with mechlorethamine gel (NM)

Corneas were allowed to recover and stabilize for 24 h before inducing NM injury. Mechlorethamine 0.016% gel (brand name Valchlor^®^, Helsinn Therapeutics, USA) was applied to the corneas as follows: ∼ 8 mg of 0.016% mechlorethamine gel was added to a 3 mm filter paper disk (Whatman^®^, USA), placed on the cornea, and incubated for 5 or 15 minutes at 37°C / 5% CO_2_. Controls were: i) unwounded corneas with no treatment; ii) corneas treated with filter paper only (FP). Then, corneas were rinsed three times with PBS and fixed immediately after wounding with 10% (w/v) paraformaldehyde (PFA, Sigma-Aldrich, Cat No. P6148, Germany) and 1% (v/v) glutaraldehyde (Sigma-Aldrich, USA) solution in PBS.

### 2.3 Histology

After fixing, corneas were paraffin-embedded and sectioned into 10 µm sections, then mounted on glass slides. Sections were stained with hematoxylin and eosin (H&E). Images of the cross-sections were taken using an Olympus microscope (Olympus BX43) and cellSens Dimension software (Olympus). Images were taken at magnifications of 4x, 10x, and 20x to observe any structural changes in epithelial thickness, epithelial loss, epithelial-stroma separation, keratocyte cells, and inflammation cell count as a consequence of mechlorethamine exposure. At least 3 corneas were imaged per condition. ImageJ (version 1.53e, National Institutes of Health, USA) was used for the measurements.

Epithelium thickness was determined by calculating the average thickness of at least five separate measurements in the wounded area per sample. The percentage of epithelium-stroma separation was calculated as = (length of total epithelial separation ÷ entire cornea length) × 100.

For Keratocyte and inflammatory cell count, we adapted the methodology used by Goswami et al. ^(6)^: the number of keratocytes and inflammatory cells was estimated from 3 different stromal areas (1 mm^2^ each), in the injury site from each cornea. A cell in the stroma with a flat nucleus was classified as a keratocyte, whereas a round nucleus was indicated an inflammatory cell.

### 2.4 Immunohistochemistry for COX-2 and FN1

IHC was performed to visualize the expression of cyclooxygenase-2 (COX-2) and fibronectin-1 (FN1) in response to mechlorethamine gel injury. Paraffin-embedded tissue slides were deparaffinized in xylene and rehydrated. Antigen retrieval was conducted in 1x citrate buffer (Cell Signaling Technology, Cat No. 14746, USA) at 98°C for 30 seconds, followed by 10 minutes at 90°C using a pressure cooker. Slides were incubated with freshly prepared 3% hydrogen peroxide for 10 minutes, then blocked with Tris buffered saline with Tween® 20 (TBST) containing 5% normal goat serum and 2.5% bovine serum albumin at room temperature for 1 hour. After blocking, slides were incubated with primary antibodies (rabbit anti-COX2 (1:300, Cell Signaling Technology, Cat No. 12282, USA) and rabbit anti-FN1 (1:100, Cell Signaling Technology, Cat No. 26836, USA)) diluted in SignalStain® Antibody Diluent (Cell Signaling Technology, USA) overnight at 4°C. On the following day, slides were washed with TBST and incubated with HRP-conjugated secondary antibody (SignalStain® Boost, HRP, Rabbit, Cell Signaling Technology, Cat No. 8114, USA) for 30 minutes at room temperature. Chromogenic detection was performed using the Epredia™ DAB Quanto Detection System (Fisher Scientific, Cat No. TA125QHDX, USA) for 3 minutes. Tissue sections were imaged under a microscope, capturing 3-7 sequential 10x images for quantification using Olympus cellSens Dimension software. COX-2- or FN1-positive cells were quantified and averaged per sample as the percentage of positive area per total tissue area or as the number of positive objects (including cells and extracellular matrix components) per mm^2^ of epithelial and stromal area.

### 2.5 Statistical analysis

One-way analysis of variance (ANOVA) with multiple comparisons (Kruskal-Wallis test) or two-tailed unpaired Student’s t-test or Mann-Whitney test were performed accordingly to identify statistical differences between the controls and test groups, using GraphPad Prism [version 10.1.2]. p< 0.05 was considered significant.

## 3. Results

We developed an NM-induced cornea injury model using *ex vivo* porcine corneas and 0.016% mechlorethamine gel. Figure 1.A shows a diagram of the lateral view of the unwounded organ culture of the porcine cornea set up. The top view of a healthy, clear cornea is seen in Figure 1.B. Following application of mechlorethamine to the cornea, an opaque area was evident at the place where NM was applied (Figure 1.C and Figure 1.D).

Corneas were fixed immediately after wounding (denominated 0 h hereafter) and H&E staining was performed on healthy, unwounded corneas (control), corneas with filter paper only (FP control), and corneas exposed to mechlorethamine gel for 5 or 15 minutes (NM 5 min and NM 15 min respectively) to observe the epithelial histopathology post-injury. Healthy epithelium was seen in unwounded and FP control corneas (Figure 2.A and 2.B respectively); whereas epithelial loss (yellow arrowhead) and epithelial thinning (red arrowhead) were observed in corneas exposed to mechlorethamine gel for 5 minutes (Figure 2.C).

**Figure 2.**
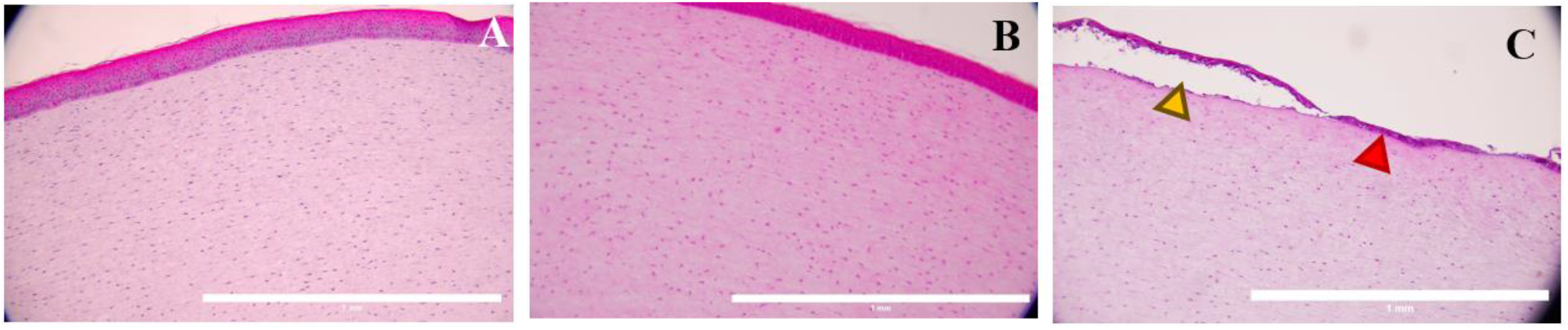
Corneal exposure to mechlorethamine gel for 5 min induces de-epithelialization typically seen in nitrogen mustard (NM)-induced corneal injury. A. Corneal model with healthy looking epithelium and stroma. B. Cornea incubated with filter paper (FP) disk, alone, for 5 min. The cornea shows a healthy epithelium and stroma at 0 h post-exposure. C. Cornea exposed to mechlorethamine gel for 5 min, showing epithelial loss (yellow arrowhead) and thinning (red arrowhead) at 0 h post-exposure. H&E staining. N=3-5. Magnification 10x. Scale bars = 1 mm.

Additionally, unwounded and FP control corneas exhibited healthy epithelium (Figure 3.A and 3.B respectively) when incubated for 15 min. On the other hand, epithelial loss (yellow arrowheads) and epithelium-stroma separation (red arrowheads) in corneas exposed to mechlorethamine gel for 15 minutes is shown in Figure 3.C. This epithelial histopathology is characteristic of NM corneal injuries.

**Figure 3.**
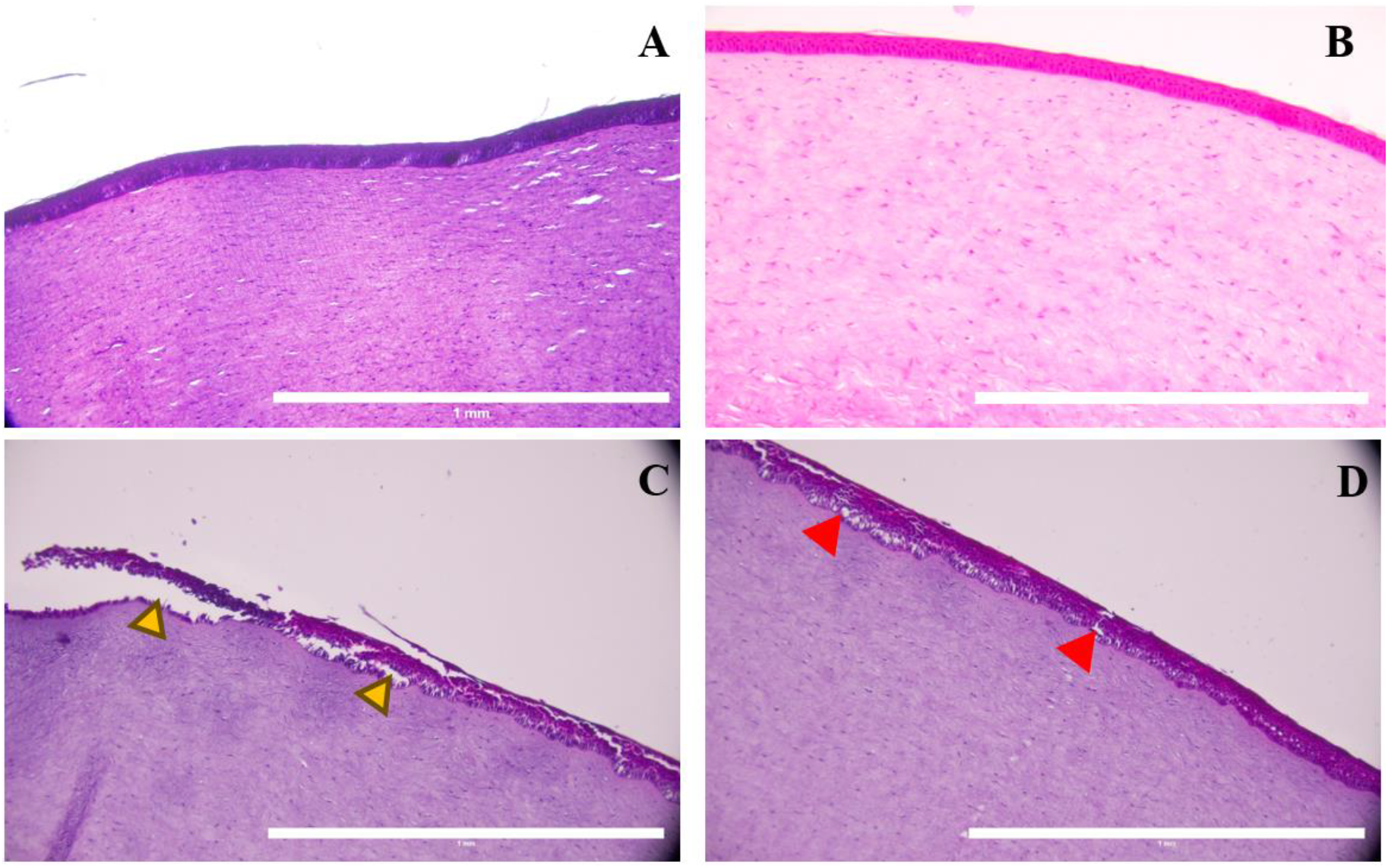
Corneal exposure to mechlorethamine gel for 15 min induces de-epithelialization and epithelium-stroma separation typically seen in nitrogen mustard (NM)-induced corneal injury. A. Corneal model with healthy looking epithelium and stroma. B. Cornea incubated with FP disk alone, for 15 min. The cornea showed a normal epithelium and stroma. C. Cornea exposed to mechlorethamine gel for 15 min, showing epithelial loss (yellow arrowheads). D. Cornea exposed to mechlorethamine gel for 15 min, showing epithelial-stroma separation (red arrowheads) at 0 h post-exposure. H&E staining. Magnification 10x. N=3-4. Scale bars = 1 mm.

We quantified epithelium thickness, epithelium-stroma separation, and keratocyte and inflammation cell counts (6,10,14,29,30) and found that average epithelium thickness was significantly reduced in both NM 5 and 15 min, from healthy control 0.062 ± 0.011 mm to 0.043 ± 0.017 mm and 0.032 ± 0.016 mm respectively (Figure 4.A). Further analysis showed that epithelium-stroma separation (Figure 4.B) was significantly higher in corneas exposed 5 and 15 min to NM (15.8% and 27% respectively) compared to healthy control (2.25%).

**Figure 4.**
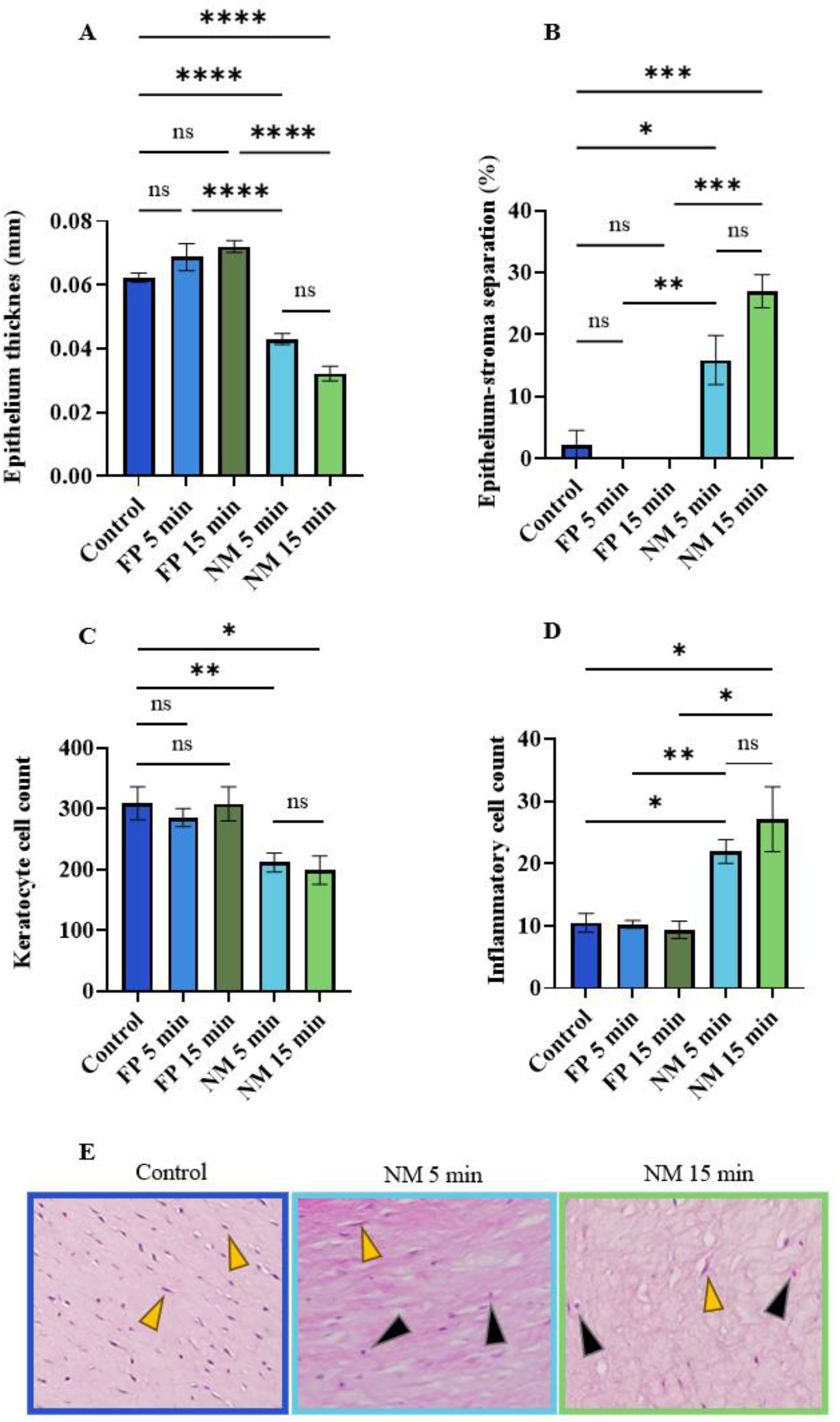
Mechlorethamine gel causes epithelium thinning, epithelium-stroma separation, a decreased keratocyte cell count and an increase in inflammatory cells in the stroma. A) Epithelium thickness decreased, and B) the percentage of epithelium-stroma separation increased after NM exposure. C) Keratocyte cells (yellow arrowheads in panel E) count decreased after NM exposure whereas D) inflammatory cells (black arrowhead in panel E) count increased. Data presented as Mean ± SEM. ANOVA with Kruskal-Wallis test and student’s t-test. ^*^p <0.05, ^**^p< 0.01,^***^p<0.001,^****^ p<0.0001. ns= no significant. N=3-5.

Keratocyte cell counts decreased significantly after NM exposure (average 211 ± 46 and 199 ± 57 keratocytes for 5 and 15 min respectively) in comparison to healthy control (average 308 ± 60 cells). Nevertheless, there was no significant difference in keratocyte numbers in samples exposed to 5 or 15 min to NM (Figure 4.C). On the other hand, the number of inflammatory cells increased significantly after NM injury (Figure 4.D) suggesting that the inflammatory response started immediately after wounding. Also, as in keratocyte cell count, the number of inflammatory cells in the stroma, below the wounded area, was not different in samples injured to NM for 5 or 15 min (average 21 ± 5 and 27 ± 13 respectively).

We stained for inflammation marker COX-2, and fibrosis marker FN1 to evaluate the inflammatory and fibrosis response after mechlorethamine gel injury. IHC showed that both COX-2 and FN1 were highly expressed in mechlorethamine gel injured corneas, for both 5- and 15-min exposure times, compared to unwounded and FP controls (Figure 5). This suggests that the inflammation and fibrotic responses started immediately after exposure.

**Figure 5.**
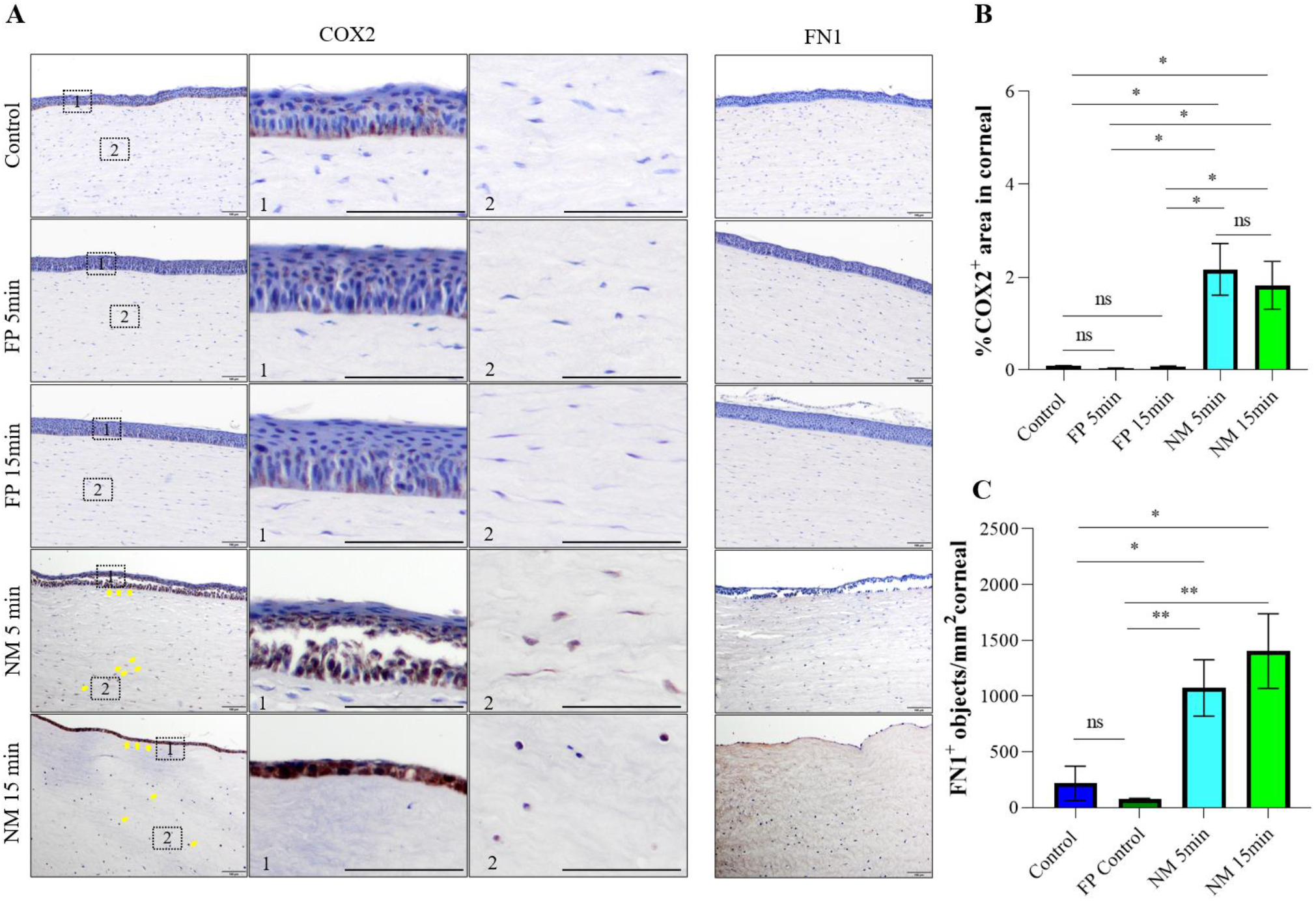
Mechlorethamine gel-induced production of COX-2 and FN1 (inflammatory and fibrotic markers, respectively) typically seen in NM corneal injury models. Porcine corneas were injured with mechlorethamine gel for 5 and 15 min, and then immediately fixed. IHC showed expression of COX-2 and FN1 in both NM injured corneas. Representative images (A) and quantification (B and C) of immunohistochemistry staining of COX2- or FN1-positive cells in NM-induced corneal injury and controls. High magnification frames on the right of the COX-2 panel show the distribution of COX-2 positive cells in epithelium and stroma of NM-treated groups. Scale bars = 100µm. Yellow arrows point to the COX2-positive keratinocytes in disrupted epithelium and stroma. Data is representative of at least 3 independent experiments with 2∼4 samples in each group were qualified using unpaired t-test or Mann-Whitney test for statistics. ^*^p <0.05, ^**^p< 0.01, ns= no significant.

## 4. Discussion

Here, we describe a safe, novel, and easy NM-induced corneal injury model, where we used mechlorethamine gel to induce an NM injury. We observed histopathology typical to that reported in well-established mustard models used by Gordon et al., Joseph et al., DeSantis-Rodrigues et al., Goswami et al., Tewari-Singh et al., Ruff et al., Charkoftaki et al., and Mishra et al., ^(6,9,10,13,14,18–20,30–33)^. The use of 0.016% mechlorethamine gel to emulate SM ocular injuries in *ex vivo* porcine cornea models is an innovative and safe method to study the injury mechanism behind SM and NM ocular exposure, as handling risks are minimized. We used porcine corneas in this model, which have the advantage that they are biologically similar to human corneas, are cost-effective, readily available, and follow the 3R’s principle ^(24–26)^. Therefore, this model will be readily available to all research laboratories, without the need for specialized facilities. Additionally, it may be the steppingstone by which novel therapeutics are developed to counteract the effects of SM.

The cornea transmits 75% of the light to the lens due to its transparency and protects the interior of the eye from any contaminants ^(34)^. Injuries and various corneal diseases may cause vision loss ^(34)^. For this reason, cornea wound healing is crucial for maintaining the integrity and functionality of the eye. Chemicals are one of the primary causes of ocular injury, affecting any or all parts of its structure. Chemical injuries may be irreversible and may continue to have long-term effects ^(35)^. SM is the most abundant alkylating agent, and it still represents a significant threat to soldiers and the civilian population due to its ease of fabrication, stockpiling, and deployment ^(1)^. Depending on SM exposure, the symptoms and treatments differ. Nevertheless, even minimum exposure to SM causes devastating injuries to the eye ^(1)^. Due to SM’s highly hazardous nature, it is very difficult to use in the laboratory, as specialized facilities, rigorous protective equipment, and permits are needed. Therefore, its analog NM has been used instead ^(13,20)^. NM exposure to the cornea causes decreased cell viability, cell death, separation of the epithelium from the stroma, neovascularization, formation of fibrotic tissue, changes in corneal and epithelial thickness, and epithelial degradation ^(14,18,20,29,36)^, and is an alternative model for laboratory research of SM-injuries. However, the use of NM also has significant limitations, since it remains a very hazardous material that requires specific protective equipment. Those limitations prevent most research laboratories from studying mustard-injuries. Consequently, models to study mustard injuries are not available to regular ocular research laboratories, hindering the development of therapeutics for these devastating injuries.

In this study, we used mechlorethamine gel to induce an NM injury in the cornea. The use of this gel does not have the health and safety risks to researchers, as mechlorethamine gel can be used by patients safely and conveniently at home. Even though the dosage of NM used herein was relatively low (0.016%), the onset of the injury was immediate, as we observed corneal opacification very soon after application (Figure 1.C). We found that mechlorethamine gel exposure (for both 5 and 15 min) causes epithelium thinning, epithelium stroma separation, decreased keratocyte count, and increased inflammatory cell count. These histopathological features are typical in well-established SM- and NM-models from Goswami et al., Tewari-Singh et al., DeSantis-Rodrigues et al., Gordon et al., Milhorn et al., Banin et al., Mishra et al., and Ruff et al., ^(9,10,12–14,18,19,29,32,33,36,37)^. Furthermore, the validity of our model is reinforced by Soleimani et al. ^(7)^ who stated that loosening of the epithelium is a key lesion of MSK; and by Kanavi et al., ^(38)^ who studied the histopathologic characteristics in patients with chronic and delayed MGK, finding keratocyte loss and irregularities in epithelium thickness.

Moreover, toxic effects of SM include edema, irritation, photophobia, corneal opacity (due to corneal scarring), and inflammation ^(4,6,8,13)^. We showed here that corneas injured with mechlorethamine gel for 5 or 15 min, expressed cyclooxygenase-2 (COX-2) and fibronectin 1 (FN1) immediately after exposure. These results suggest that the response to NM injury is rapid, and that both exposure times may be used to further study the effects of NM in the cornea. COX-2 has been reported as a critical mediator in NM and SM-induced inflammation in the cornea ^(13,39,40)^. Goswami et al., Tewari-Singh et al., and Mishra et al., showed an increase in COX-2 in both their SM- and NM-corneal models ^(6,10,13,14,19,29,33)^. Additionally, FN1, a key molecule related to scar formation in the cornea, is expressed during early phases of wound healing ^(41,42)^. Joseph et al. showed altered expression of FN1 in rabbit corneas after SM injury ^(30)^. Research involving *in vivo* ocular experiments established structural alterations, inflammation, neovascularization, and opacity when eyes were exposed to SM for less than 4 minutes ^(43–46)^.

Interestingly, there is a major gap in reporting the effects of NM or SM in the cornea immediately after exposure. Most of the studies describe histopathology after 24 h of mustard exposure, with no mention of any effects observed earlier than that time. One reason for this is that, for the safety of the researchers, initial evaluations were done 24 h post injury to minimize or avoid the risk of SM vapor emission ^(46)^. However, it is not clear why, in models that used liquid NM, the injury effects were not evaluated soon after exposure. To the best of our knowledge, Charkofttaki et al., assessed corneal structure post NM injury but found no morphological changes 3 h after NM exposure ^(20)^. Thus, the advantage of using the safe NM corneal injury model presented herein is that researchers can evaluate safely the effects of NM exposure immediately after injury, facilitating the identification of early (within 24 h post-injury) or possibly new molecules and mechanisms that might help to increase our understanding of mustard ocular injuries as well to identify novel therapeutics. Nevertheless, the limitations of using this safe model with mechlorethamine gel are that only 0.016% or lower concentrations of NM can be used, making it difficult to study the effects of higher concentrations; also, it does not mimic how eyes are injured in the battlefield, as the injury is only localized in a specific area of the cornea. The size of the injury might be overcome by using a bigger filter paper disk. However, further evaluation is needed to identify if longer exposure times would increase the severity of the NM injury. Future use of this model *in vivo* would allow for the assessment of long-term effects of NM toxicity, and for the investigation of novel therapies for mustard corneal exposure.

In conclusion, we report a novel, safe, and very practical model of NM-induced corneal injury that showed the characteristic histopathology, and expression of inflammation (COX-2) and fibrotic (FN1) markers of NM injury in the cornea, which are consistent with other well-established SM- and NM-corneal injury models. The use of mechlorethamine 0.016% gel to mimic SM ocular injuries in *ex vivo* porcine corneas can be adapted by any laboratories as a safe method to study the mechanisms behind mustard ocular exposure and injury, without the need for any special facilities or equipment. It is especially suitable to study the very early pathological features of mustard injuries within minutes to hours after exposure. This will shed light on new knowledge that would help us to increase our understanding of the mechanisms and effects of mustard ocular injuries while investigating novel therapeutics.

## 5. Acknowledgments

Dr. Brian Reid for his insights on this project and for proofreading this manuscript. Wen Min for her technical support.

## References

1. Morad Y, Banin E, Averbukh E, Berenshtein E, Obolensky A, Chevion M. Treatment of ocular tissues exposed to nitrogen mustard: beneficial effect of zinc desferrioxamine combined with steroids. Invest Ophthalmol Vis Sci. 2005;46(5):1640–6.

2. Graham JS, Schoneboom BA. Historical perspective on effects and treatment of sulfur mustard injuries. Chem Biol Interact. 2013;206(3):512–22.

3. Lombardo C. National Public Radio. 2019. More Than 300 Chemical Attacks Launched During Syrian Civil War, Study Says. Available from: https://www.npr.org/2019/02/17/695545252/more-than-300-chemical-attackslaunched-during-syrian-civil-war-study-says

4. Mishra N, Agarwal R. Research models of sulfur mustard-and nitrogen mustard-induced ocular injuries and potential therapeutics. Exp Eye Res. 2022;223:109209.

5. Araj H, Tseng H, Yeung DT. Supporting discovery and development of medical countermeasures for chemical injury to eye and skin. Exp Eye Res. 2022;221:109156.

6. Goswami DG, Mishra N, Kant R, Agarwal C, Croutch CR, Enzenauer RW, et al. Pathophysiology and inflammatory biomarkers of sulfur mustard-induced corneal injury in rabbits. PLoS One. 2021;16(10):e0258503.

7. Soleimani M, Momenaei B, Baradaran-Rafii A, Cheraqpour K, An S, Ashraf MJ, et al. Mustard Gas–Induced Ocular Surface Disorders: An Update on the Pathogenesis, Clinical Manifestations, and Management. Cornea. 2022;42(6):776–86.

8. Fuchs A, Giuliano EA, Sinha NR, Mohan RR. Ocular toxicity of mustard gas: A concise review. Toxicol Lett. 2021;343:21–7.

9. Ruff AL, Jarecke AJ, Hilber DJ, Rothwell CC, Beach SL, Dillman III JF. Development of a mouse model for sulfur mustard-induced ocular injury and long-term clinical analysis of injury progression. Cutan Ocul Toxicol. 2013;32(2):140–9.

10. Goswami DG, Mishra N, Kant R, Agarwal C, Ammar DA, Petrash JM, et al. Effect of dexamethasone treatment at variable therapeutic windows in reversing nitrogen mustard-induced corneal injuries in rabbit ocular in vivo model. Toxicol Appl Pharmacol. 2022;437:115904.

11. Rafati-Rahimzadeh M, Rafati-Rahimzadeh M, Kazemi S, Moghadamnia AA. Therapeutic options to treat mustard gas poisoning–Review. Caspian J Intern Med. 2019;10(3):241.

12. Milhorn D, Hamilton T, Nelson M, McNutt P. Progression of ocular sulfur mustard injury: development of a model system. Ann N Y Acad Sci. 2010;1194(1):72–80.

13. Goswami DG, Tewari-Singh N, Dhar D, Kumar D, Agarwal C, Ammar DA, et al. Nitrogen mustard-induced corneal injury involves DNA damage and pathways related to inflammation, epithelial-stromal separation and neovascularization. Cornea. 2016;35(2):257.

14. Goswami DG, Kant R, Ammar DA, Kumar D, Enzenauer RW, Petrash JM, et al. Acute corneal injury in rabbits following nitrogen mustard ocular exposure. Exp Mol Pathol. 2019;110:104275.

15. Panahi Y, Roshandel D, Sadoughi M, Ghanei M, Sahebkar A. Sulfur mustard-induced ocular injuries: update on mechanisms and management. Curr Pharm Des. 2017;23(11):1589–97.

16. Thompson VR, DeCaprio AP. Covalent adduction of nitrogen mustards to model protein nucleophiles. Chem Res Toxicol. 2013;26(8):1263–71.

17. Wattana M, Bey T. Mustard gas or sulfur mustard: an old chemical agent as a new terrorist threat. Prehosp Disaster Med. 2009;24(1):19–29.

18. Gordon MK, DeSantis A, Deshmukh M, Lacey CJ, Hahn RA, Beloni J, et al. Doxycycline hydrogels as a potential therapy for ocular vesicant injury. Journal of ocular pharmacology and therapeutics. 2010;26(5):407–19.

19. Tewari-Singh N, Jain AK, Inturi S, Ammar DA, Agarwal C, Tyagi P, et al. Silibinin, dexamethasone, and doxycycline as potential therapeutic agents for treating vesicant-inflicted ocular injuries. Toxicol Appl Pharmacol. 2012;264(1):23–31.

20. Charkoftaki G, Jester J V, Thompson DC, Vasiliou V. Nitrogen mustard-induced corneal injury involves the sphingomyelin-ceramide pathway. Ocul Surf. 2018;16(1):154–62.

21. Merck KGaA [Internet]. 2022 [cited 2022 Apr 25]. Mechlorethamine hydrochloride. Available from: https://www.sigmaaldrich.com/US/en/product/aldrich/122564

22. Meng W, Sun M, Xu Q, Cen J, Cao Y, Li Z, et al. Development of a series of fluorescent probes for the early diagnostic imaging of sulfur mustard poisoning. ACS Sens. 2019;4(10):2794–801.

23. Agarwal P, Rupenthal ID. In vitro and ex vivo corneal penetration and absorption models. Drug Deliv Transl Res. 2016;6(6):634–47.

24. Seyed-Safi AG, Daniels JT. A validated porcine corneal organ culture model to study the limbal response to corneal epithelial injury. Exp Eye Res. 2020;197:108063.

25. Okurowska K, Roy S, Thokala P, Partridge L, Garg P, MacNeil S, et al. Establishing a porcine ex vivo cornea model for studying drug treatments against bacterial keratitis. JoVE (Journal of Visualized Experiments). 2020;(159):e61156.

26. Menduni F, Davies LN, Madrid-Costa D, Fratini A, Wolffsohn JS. Characterisation of the porcine eyeball as an in-vitro model for dry eye. Contact Lens and Anterior Eye. 2018;41(1):13–7.

27. Helsinn Therapeutics Inc. VALCHLOR. 2024.

28. Castro N, Gillespie SR, Bernstein AM. Ex vivo corneal organ culture model for wound healing studies. JoVE (Journal of Visualized Experiments). 2019;(144):e58562.

29. Goswami DG, Kant R, Tewari-Singh N, Agarwal R. Efficacy of anti-inflammatory, antibiotic and pleiotropic agents in reversing nitrogen mustard-induced injury in ex vivo cultured rabbit cornea. Toxicol Lett. 2018;293:127–32.

30. Joseph LB, Gordon MK, Zhou P, Hahn RA, Lababidi H, Croutch CR, et al. Sulfur mustard corneal injury is associated with alterations in the epithelial basement membrane and stromal extracellular matrix. Exp Mol Pathol. 2022;128:104807.

31. DeSantis-Rodrigues A, Chang YC, Hahn RA, Po IP, Zhou P, Lacey CJ, et al. ADAM17 inhibitors attenuate corneal epithelial detachment induced by mustard exposure. Invest Ophthalmol Vis Sci. 2016;57(4):1687–98.

32. Mishra N, Kant R, Kandhari K, Ammar DA, Tewari-Singh N, Pantcheva MB, et al. Nitrogen mustard-induced ex vivo human cornea injury model and therapeutic intervention by dexamethasone. Journal of Pharmacology and Experimental Therapeutics. 2024;388(2):484–94.

33. Mishra N, Kant R, Kandhari K, Tewari-Singh N, Anantharam P, Croutch CR, et al. Establishing a Dexamethasone Treatment Regimen To Alleviate Sulfur Mustard– Induced Corneal Injuries in a Rabbit Model. Journal of Pharmacology and Experimental Therapeutics. 2024;388(2):469–83.

34. Mahdavi SS, Abdekhodaie MJ, Mashayekhan S, Baradaran-Rafii A, Djalilian AR. Bioengineering Approaches for Corneal Regenerative Medicine. Tissue Eng Regen Med. 2020;17(5):567–93.

35. Ashtiani HRA, Garmroodi ARN, Hazrati E. A Review of Management Methods and Modern Treatments for Chemical Wounds. Journal of Archives in Military Medicine. 2021;9(1):e112029.

36. Banin E, Morad Y, Berenshtein E, Obolensky A, Yahalom C, Goldich J, et al. Injury induced by chemical warfare agents: characterization and treatment of ocular tissues exposed to nitrogen mustard. Invest Ophthalmol Vis Sci. 2003;44(7):2966–72.

37. DeSantis-Rodrigues A, Hahn RA, Zhou P, Babin M, Svoboda KKH, Chang Y, et al. SM1997 downregulates mustard-induced enzymes that disrupt corneal epithelial attachment. Anat Rec. 2021;304(9):1974–83.

38. Kanavi MR, Javadi A, Javadi MA. Chronic and delayed mustard gas keratopathy: a histopathologic and immunohistochemical study. Eur J Ophthalmol. 2010;20(5):839– 43.

39. Kumar D, Tewari-Singh N, Agarwal C, Jain AK, Inturi S, Kant R, et al. Nitrogen mustard exposure of murine skin induces DNA damage, oxidative stress and activation of MAPK/Akt-AP1 pathway leading to induction of inflammatory and proteolytic mediators. Toxicol Lett. 2015;235(3):161–71.

40. Shakarjian MP, Heck DE, Gray JP, Sinko PJ, Gordon MK, Casillas RP, et al. Mechanisms mediating the vesicant actions of sulfur mustard after cutaneous exposure. Toxicological sciences. 2010;114(1):5–19.

41. Chang Y, Wu XY. The role of c-jun N-terminal kinases 1/2 in transforming growth factor β1-induced expression of connective tissue growth factor and scar formation in the cornea. Journal of International Medical Research. 2009;37(3):727–36.

42. Barbariga M, Vallone F, Mosca E, Bignami F, Magagnotti C, Fonteyne P, et al. The role of extracellular matrix in mouse and human corneal neovascularization. Sci Rep. 2019;9(1):14272.

43. Kadar T, Dachir S, Cohen L, Sahar R, Fishbine E, Cohen M, et al. Ocular injuries following sulfur mustard exposure—pathological mechanism and potential therapy. Toxicology. 2009;263(1):59–69.

44. McNutt P, Hamilton T, Nelson M, Adkins A, Swartz A, Lawrence R, et al. Pathogenesis of acute and delayed corneal lesions after ocular exposure to sulfur mustard vapor. Cornea. 2012;31(3):280–90.

45. Kadar T, Turetz J, Fishbine E, Sahar R, Chapman S, Amir A. Characterization of acute and delayed ocular lesions induced by sulfur mustard in rabbits. Curr Eye Res. 2001;22(1):42–53.

46. McNutt PM, Kelly KEM, Altvater AC, Nelson MR, Lyman ME, O’Brien S, et al. Dose-dependent emergence of acute and recurrent corneal lesions in sulfur mustard-exposed rabbit eyes. Toxicol Lett. 2021;341:33–42.

