## Supplemental Table 1 for "A Practical and Safe Model of Nitrogen Mustard Injury in Cornea"

Table S1. Reagents used to prepare culture medium for ex vivo corneal culture.

| Reagent | Final concentration | Cat No. | Manufacturer | Country |
| --- | --- | --- | --- | --- |
| DMEM/F-12 | NA | 11330032 | Life Technologies | USA |
| RPMI 1640 vitamins solution | 1% v/v | R7256 | Sigma-Aldrich | Germany |
| ITS liquid media supplement | 1% v/v | I3146 | Sigma-Aldrich | Germany |
| L-Glutathione reduced | 1 µg/mL | G6013 | Sigma-Aldrich | Germany |
| L-Glutamine | 1% | 25030081 | Thermo Fisher Scientific | UK |
| MEM non-essential amino acids solution | 1% | 11140050 | Thermo Fisher Scientific | USA |
| Sodium Pyruvate | 1 mM | 11360070 | Thermo Fisher Scientific | USA |
| Antibiotic Antimycotic solution (ABAM) | 1% | A5955 | Sigma-Aldrich | Germany |
| Gentamicin | 50 µg/mL | G1397 | Sigma-Aldrich | Germany |
| 2-O-s-D-Glucopyranosyl-L-ascorbic acid | 1 mM | SMB00390 | Sigma-Aldrich | Germany |
